# The Development of 2-stage Microfermentation Protocols for High Throughput Cell Factory Evaluations

**DOI:** 10.1101/2022.02.25.481916

**Authors:** Shuai Li, Zhixia Ye, Eirik A. Moreb, Romel Menacho-Melgar, Michael D. Lynch

## Abstract

Cell based factories can be engineered to produce a wide variety of products. Advances in DNA synthesis and genome editing have greatly simplified the design and construction of these factories. It has never been easier to generate hundreds or even thousands of cell factory strain variants for evaluation. These advances have amplified the need for standardized, higher throughput means of evaluating these designs. Toward this goal, we have previously reported the development of engineered *E. coli* strains and associated 2-stage production processes to simplify and standardize strain engineering, evaluation and scale up. This approach relies on decoupling growth (stage 1), from production, which occurs in stationary phase (stage 2). Phosphate depletion is used as the trigger to stop growth as well as induce heterologous expression. Here, we describe in detail the development of optimal protocols used for the evaluation of engineered *E. coli* strains in 2-stage microfermentations. These protocols are readily adaptable to the evaluation of strains producing a wide variety of protein as well as small molecule products. Additionally, the development approach described is adaptable to additional cellular hosts, as well as other 2-stage processes with various additional triggers.

## Introduction

The past decade has seen significant advances in synthetic biology, in particular advances in our ability to write DNA and edit genomes.^1,2^ It has never been easier to engineer living systems. Advances in synthetic biology in combination with advances in protein and enzyme engineering, expression and purification and metabolic engineering, have enabled numerous advanced cell factories for the production of proteins as well as metabolites or chemicals.^3–5^ Among these cell factories, *E. coli* remains a workhorse microbe for the production of numerous products, including both proteins (including therapeutics) and small molecules.^3,6–8^ *E. coli* is well-studied and has well-developed high cell density culturing techniques, as well as established genome, metabolic and protein engineering tools. ^9,10^

Advanced tools for genetic manipulation have made it easier to generate a wide diversity of strain designs, necessitating advances in higher throughput methods for testing or evaluating these designs. ^11^ This need has led to the development of numerous “scale” down systems to enable strain evaluation or process development at smaller scales in higher throughput.^12–20^ While many of these “microbioreactor” systems offer significant advances when compared to traditional shake flask or microtiter plate cultivation methods, they are not readily accessible as they can be expensive and require significant adaptation to a given set of microbial strain variants and/or target product. We have recently reported the development of a methodology enabling 2-stage production of both proteins and small molecules.^21–23^ This approach leverages engineered strains of *E. coli* and standardized (product independent) methods for 2-stage production in instrumented bioreactors as well as microtiter plates. Importantly, this methodology enables predictability across scales from microtiter plates to instrumented reactors. ^22,24^ This enables reliable scale up of a strain or strain variant identified using the microfermentation protocol.

Importantly, as illustrated in Figure 1, we have developed two microfermenation protocols for the rapid and standardized evaluation of *E. coli* cell factories. Both of these 2-stage protocols rely on phosphate depletion to stop cell growth and induce a production stage. The first, Figure 1a, is enabled by autoinduction media enabling a “hands off” protocol.^23,25^ The second Figure 1b, is based on an initial growth stage, followed by washing cells to induce the production phase. ^21,24^ This protocol requires more “hands on” time, but allows for the normalization of cell numbers/biomass levels and more control over the initial media. Both protocols also leverage the System Duetz plate cover sealing system for microtiter plate cultivations. This system (available from Enzyscreen) leverages reusable plate covers and clamps to enable adequate culture aeration, while minimizing evaporation.

**Figure 1:**
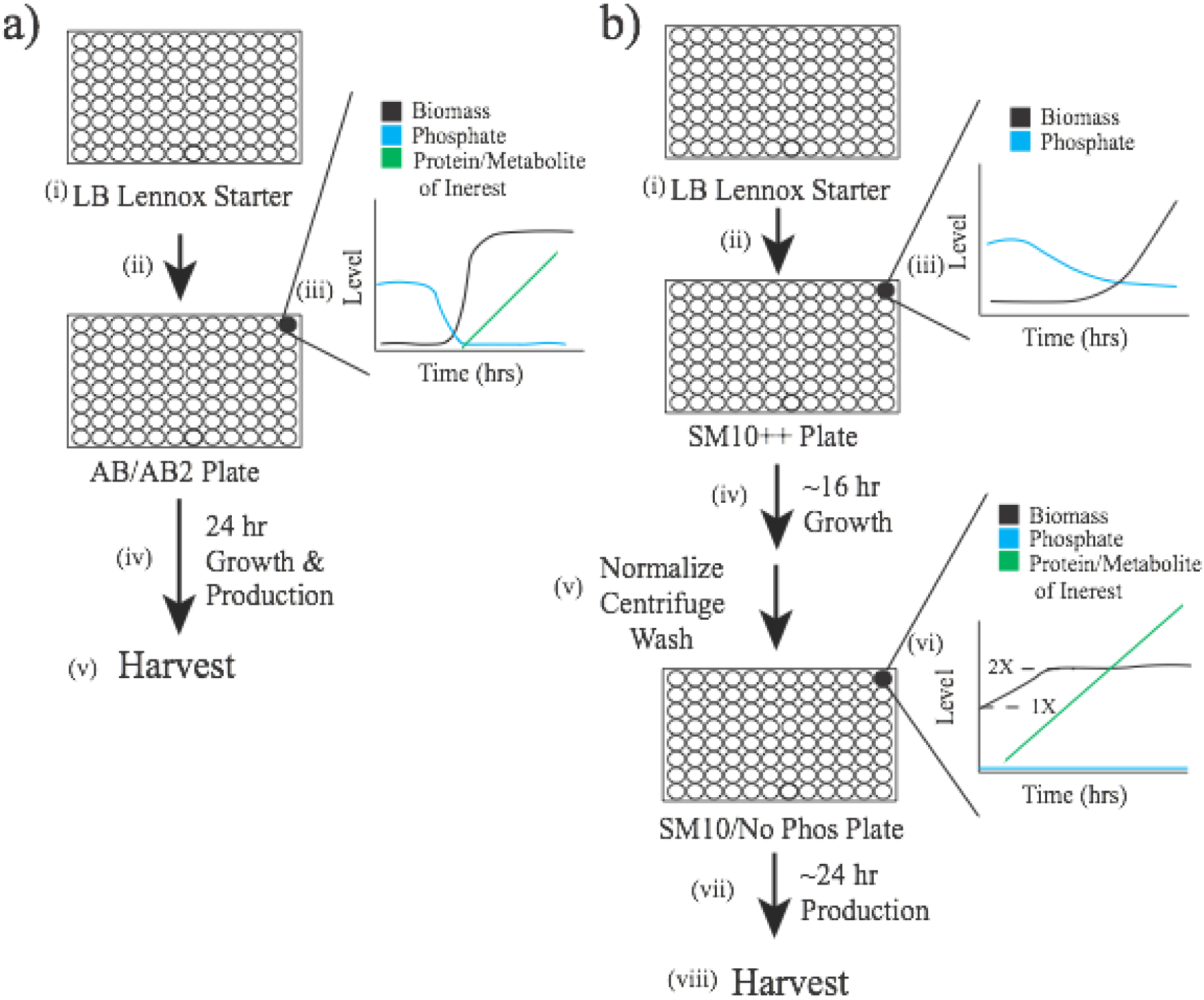
An overview of the 2-stage microfermentation protocols. a) An overview of the 2-stage autoinduction microfermentation protocol. (i) Overnight (16 hr) cultures in LB (Lennox formulation) are used to (ii) inoculate a new microtiter plate containing rich Autoinduction Broth at 1% v/v (AB or AB2). (iii) A time course of the autoinduction culture. Cells (biomass) grow and consume phosphate, which is depleted upon entry into stationary phase. Upon phosphate depletion, low phosphate inducible promoters are activated leading to expression of a target protein or metabolic pathway of interest, which in turn leads to accumulation of the target protein or metabolite. (v) After production cells or supernatants are harvested by centrifugation. b) An overview of the 2-stage wash microfermentation protocol. (i) Again overnight (16 hr) cultures in LB (Lennox formulation) are used to (ii) inoculate a new microtiter plate, in this case containing SM10++ media. SM10++ is a minimal media supplemented with yeast extract and casamino acids in order to reduce the lag of cell growth in LB. (iii) A time course of the SM10 ++ culture. Cells (biomass) grow and consume phosphate, which is not depleted. (iv) Cells are grown to mid-exponential phase (OD_600nm_ 5-10), which usually takes ∼ 16 hrs. (v) Cells are harvested by centrifugation washed with SM10 No phosphate media to remove phosphate and normalized to a target optical density (usually OD_600nm_ ∼ 1), which begins the phosphate depleted production phase. (vi) A time course of the SM10 No phosphate culture. Cells usually undergo ∼ 1 doubling, coincident with induction of low phosphate inducible promoters leading to expression of a target protein or metabolic pathway of interest, which in turn leads to accumulation of the target protein or metabolite. (v) After production cells or supernatants are harvested by centrifugation.

We have developed these 2-stage microfermentation protocols in order to enable the high throughput and reproducible evaluation of microbial strains, specifically *E. coli*. ^23–30^ While the specific protocols described herein have been optimized for 2-stage production of either proteins or metabolites upon phosphate depletion, the approach and methodology is readily adaptable to additional induction systems and microbial species. We have included, as discussed below, key considerations and results obtained during the development of the protocol, which provide i) crucial background, ii) the impact of key variables including evaporation, oxygen transfer and batch media formulation and iii) key limitations for anyone seeking to not only reproduce this specific method, but also looking to extend and/or adapt it.

### Protocol Development

#### Boundary Conditions for Small Scale Batch Cultivation

To begin, it is critical to consider the essential aspects of developing any microbial process, whether in a larger scale shake flask, instrumented bioreactor, or high throughput system such as microtiter plates. Essentially, microbial cultures need several things, a media providing a carbon source and other key nutrients for growth, pH control (such as by buffering), and adequate aeration and oxygenation (in the case of an aerobic culture). Larger intensified processes, such as those in bioreactors, enable feeding of nutrients, air and titrants, providing real time control over oxygen and pH levels. As discussed, there have been numerous systems developed for high throughput cultivation that try to mimic fed batch control available in larger bioreactors. ^12–20^ These systems are often complex, very expensive and require significant troubleshooting such as PID tuning for a given application.

When using inexpensive, standard microtiter plates, researchers are often forced to consider batch cultivation, in which all nutrients and buffers must be included at the start of the culture. Therefore it is critical to balance the expected (or targeted) microbial growth rates, product formation rates as well as final biomass and product levels with the “batch capacity” of the media. For example, you must have enough feedstock (in our case glucose) to achieve a given biomass level. Additionally, if you hope to achieve a certain level of a small molecule you need to make sure the culture can accommodate this. For example if you are expecting significant final titers of an organic acid you need to have not only enough glucose to support this production, but also enough buffering to maintain pH while the acid is accumulating. If the small molecule produced is an amino acid, buffering is less important, but having enough ammonia (a precursor to amino acids) to support a target product level is critical.

As a result of these constraints, we recommend higher cell density cultures when the product is an intracellular protein (enabling higher product levels for subsequent analysis). The autoinduction protocol (Figure 1a) results in relatively high biomass levels (particularly for microtiter plates) and is ideal for protein expression. We recommend targeting lower cell densities when the product is a small molecule. For very productive cells, a significant amount of batch glucose is needed to make the product, leaving proportionally less glucose consumption for biomass production. The 2-stage wash microfermentation protocol (Figure 1b), which normalizes cell numbers can be used, not only to control biomass levels, but also to enable better control over the media environment during the productive stationary phase. As metabolism can be sensitive to media components, this is an advantage in metabolic engineering studies. This is not to say that the autoinduction protocol cannot be leveraged in metabolic engineering programs, but we have recently demonstrated that for small molecules, performance of the 2-stage wash protocol in microtiter plates is predictive of performance in larger minimal media bioreactors, again a least for a subset of our engineered *E. coli*. ^21,22^

#### Aeration & Oxygen Transfer

Optimal oxygen transfer is essential for aerobic growth and production. A well-mixed culture is necessary for efficient oxygen supply. Optimal aeration in microtiter plates requires consideration of shaking speed as well as the orbital radius of the shaker. ^31,32^ We use shakers from Kuhner such as Model Climo-Shaker ISF4-X. It should be noted that many shakers have shaking orbits smaller than 50 mm (or different orbits). Plate fill volumes or speeds may need to be changed experimentally to ensure adequate aeration if using a different incubator. We would refer the reader to Enzyscreen’s website for options that can provide an adequate OTR (https://www.enzyscreen.com/oxygen_transfer_rates.htm) to identify alternative conditions that can meet OTR targets, when using a different orbit. Square well plate may also be used to increase OTRs to the needed levels. Alternatively, to supply anaerobic conditions, we recommend using aluminum foil or other adhesive films (such as AlumaSeal CS Sealing Films from EXCEL scientific Catalog# FSC-25) to seal the wells enabling anaerobic growth and/or production.

#### Evaporation in Microtiter Plates

As mentioned, these protocols rely on the microtiter plate covers and clamps (the Duetz System) from Enzyscreen. The covers have been designed to enable adequate aeration and oxygen transfer while minimizing the impact of evaporation, particularly at high shaking speeds.^33^ Despite the dramatic improvements observed using the Duetz System™, evaporation is still observed and we have measured this using standard flat bottom, 96 well plates. Briefly, we used the dye bromophenol blue to measure evaporation rates in simulated microfermentations. Simulated microfermentations were performed using microtiter plates filled with media (150 µl) without cells, initially containing 10 mg/L of bromophenol blue. These plates were incubated at 37°C with plate covers, shaking at 300 rpm at a 50 mm orbit. The dye is concentrated in response to evaporation. At time intervals the concentration of bromophenol blue was measured via absorbance at 590 nm, and the volume change (percent evaporation/loss) was calculated. This methodology and the associated results are given in Figure 2. As shown in Figure 2b-c, there is significantly greater evaporation rate from wells on the edge of the microtiter plate, which is a well known challenge when using microtiter plates.^33^ While this can be affected by how well the plate covers are clamped, we recommend not using the wells on the outer plate edge. Additionally, as shown in Figure 2d, the degree of evaporation increases with the time, as would be expected. After 12 hours, on average the plate had 4.6% volume loss, after 24 hours, 11.6% volume loss, after 36 hours, 19.4% volume loss, and after 48 hours, a 28.6% volume loss. This should be taken into account particularly when comparing measurements from different time points, where an increased product concentration would be expected based upon evaporation alone.

**Figure 2:**
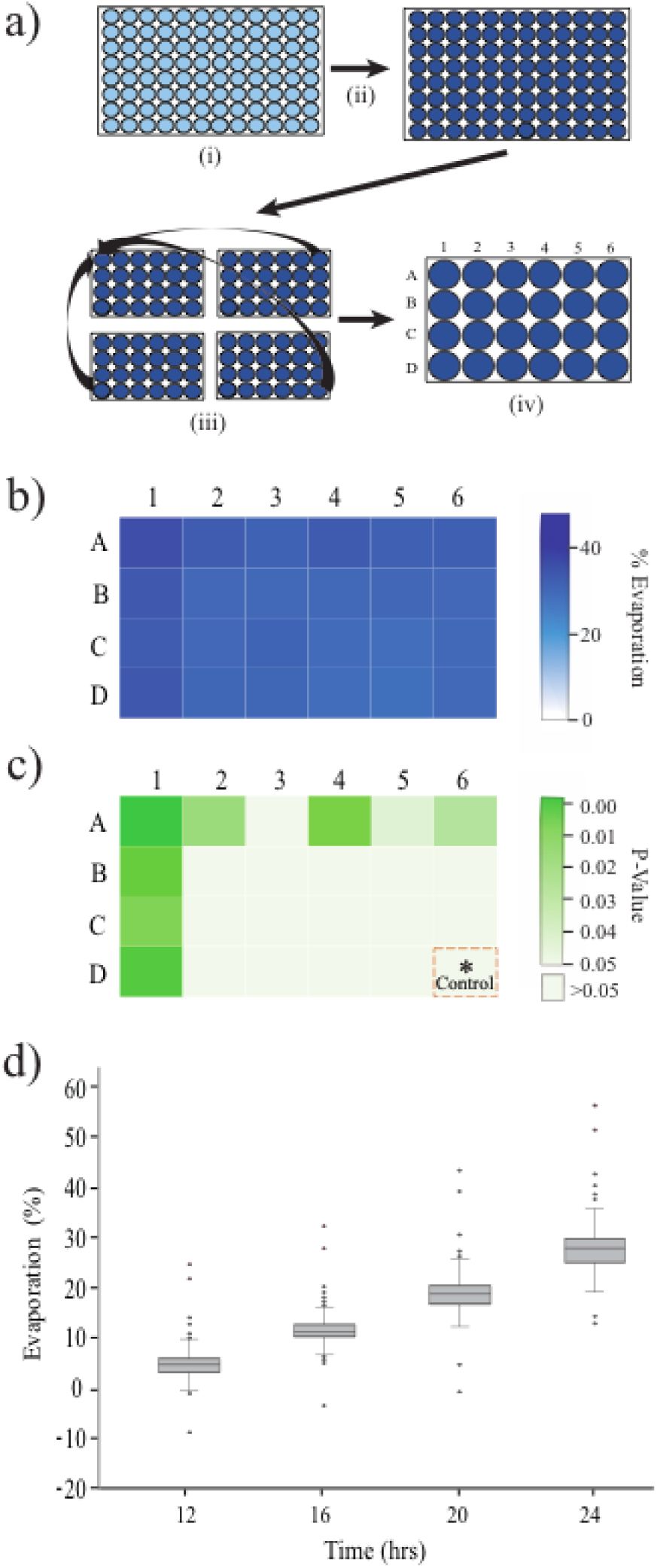
Evaporation in Micro-fermentations. a) An overview of the methodology used to measure evaporation in a simulated culture. (i) Bromophenol blue was added to the media, which was used to fill 96 well plates. (ii) The relative concentration of bromophenol blue was measured (via absorbance) as a function of time throughout a simulated microfermentation. (iii) Due to symmetry, quadrants of each plate were treated as replicates and (iv) averaged to yield a percent evaporation as a function of well position in a quadrant. b) Average percent evaporation after 48 hrs of a simulated culture as a function of well position. c) Statistical differences in evaporation as a function of position, compared to the center well. d) A time course of evaporation for 96 well simulated microfermentations. These data exclude the out edge of the microtiter plate. n = 360 for each time point.

#### Batch Media Development

Media is another critical consideration when performing high throughput microbial cultivations. As discussed above, batch media needs to be developed to provide enough feedstock and nutrients to balance biomass growth and production without requiring toxic levels. We have previously reported the approach in detail for the development of the phosphate limited autoinduction media (AB and AB2) used in the 2-stage autoinduction protocol (Figure 1a) and refer the reader to these primary sources for further information.^23,25^ AB and AB2 can be used somewhat interchangeably with respect to 2-stage protein expression, as AB2 is a streamlined and simplified version of AB. These two media have different levels of many components including trace metals which should be considered when producing metabolic products.

While we have previously reported “optimal” media used for the 2-stage wash microfermentation protocol (Figure 1b), namely SM10++ and SM10 No Phosphate, it is important to share some details of how these media were chosen, to enable the more rapid adaptation of this protocol. SM10 (Shake Flask Media 10, enabling biomass levels of 10gCDW/L) is almost identical to a minimal media developed for use in instrumented bioreactors as reported by Menacho-Melgar et al. ^25^ ie FGM10 (Fermentation Growth Media 10, enabling biomass levels of 10gCDW/). The FGM10 media formulation is similar to many previously used minimal mineral salts media containing ammonium salts, phosphate, trace metals and glucose. ^25,34,35^ One of the primary differences between FGM10 media and SM10 media is that SM10 media contains a buffer (in this case MOPS, 3-morpholinopropane-1-sulfonic acid) to control pH. Both of these media are formulated to enable *E. coli* biomass levels of ∼8-10 gCDW/L (OD_600nm_ 25-30), at which point phosphate, the limiting nutrient, is exhausted.

Unfortunately, most strains of *E*.*coli* propagated in routine complex media, such as Luria Broth, require a significant adaptation period to enable rapid growth in these minimal defined media.^36,37^As a result the direct transfer of starter cultures from complex rich media to defined media, oftentimes leads to a rapid growth period where nutrients remaining in the inocula are exhausted followed by a lag phase. The lag time (needed for media adaptation) can be significant, unpredictable and vary between strains and replicate cultures.^38–42^ In bioreactor studies, seed cultures can be performed in minimal media to allow stains to adapt prior to inoculating reactors, however this approach is not amenable to routine high throughput experimentation. Another approach is the use of a “Bridging Seed Media” which enables predictable and rapid growth while conditioning the cells to a nutrient composition closer to the defined minimal media. In order to implement this approach we developed SM10++ media which is SM10 media with the addition of small amounts of casamino acids as well as yeast extract. The impact of these nutrients on the growth of cultures directly inoculated from LB starter cultures is shown in Figure 3a and b. Briefly, 5mL, LB Lennox starter cultures were used to inoculate wells of a BioLector ™ plate (enabling real time growth monitoring) containing various media. While inoculation directly into defined minimal SM10 media results in a prolonged lag phase, the addition of casamino acids and yeast extract alone and in combination reduce the lag. SM10 ++ (“+” 2.5g/L casamino acids and “+” 2.5g/L yeast extract) was chosen as our bridging seed media due to a higher growth rate and final optical density. In this media background, the batch glucose and buffer concentrations were then also optimized as illustrated in Figure 3c-f. The goal was to maximize the batch sugar as well as buffer capacity without greatly impacting growth rates or lag times. For the current media formulations 200mM MOPS and 45g/L glucose were chosen.

**Figure 3:**
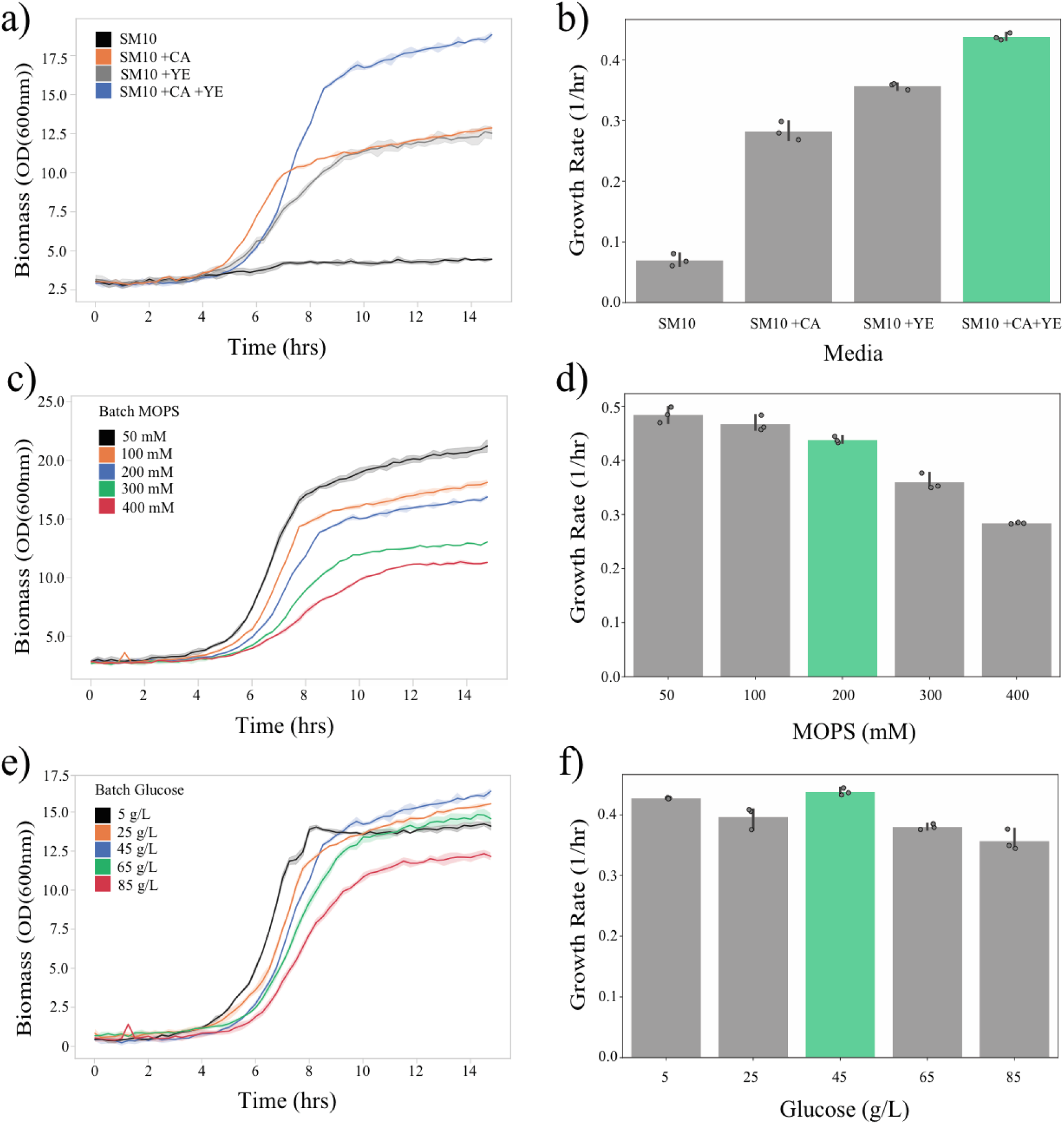
Initial SM10++ Media Development. Growth curves (a) and maximal growth rates (b) of strain DLFS_0025 with plasmids pHCKan-yibDp-GFPuv in SM10 media and SM10 media with the complex additives casamino acids (CA) and yeast extract (YE) alone and in combination. Growth curves (c) and maximal growth rates (d) of strain DLFS_0025 with plasmids pHCKan-yibDp-GFPuv in SM10++ media (with both CA and YE) with varying concentrations of MOPS buffer. Growth curves (e) and maximal growth rates (f) of strain DLFS_0025 with plasmids pHCKan-yibDp-GFPuv in SM10++ media (with both CA and YES) with varying concentrations of batch glucose. All growth curves were measured using a BioLector ™ (n=3). Abbreviations: CA - casamino acids, YE - yeast extract.

#### Key Process Variables

While the above discussion has focused on some key variables to consider in developing and performing microfermentations, other process variables can also impact performance. In the case of the autoinduction protocol (Figure 1a), there is minimal user intervention and the only real parameter that is varied is the time at which the cultures are harvested and analyzed. Results for 2-stage auto-induced microfermentations utilizing AB2 media are given in Figure 4. These results were obtained using Protocol 1. Specifically, GFPuv was expressed via a plasmid (pHCKan-GFPuv, Addgene #127078) under the control of the robust yibDp gene promoter. ^24^ These results are typical of an expected expression time course leveraging induction by phosphate depletion in *E. coli* DLF_R004 (used to generate these results) and its derivatives. Based on these results we recommend a 24 hour protocol from inoculation to harvest, as prolonging the culture does not improve protein titers. Prolonged auto induced cultures may be worth evaluating if adapting this protocol to small molecule production. Similarly, we recommend a 24 hour “production phase” when using the 2-stage wash protocol (Figure 1b). This is based on typical results such as those given in Figure 5. These results were obtained using Protocol 2, leveraging strain DLFS_0025 bearing plasmid pHCKan-GFPuv.

**Figure 4:**
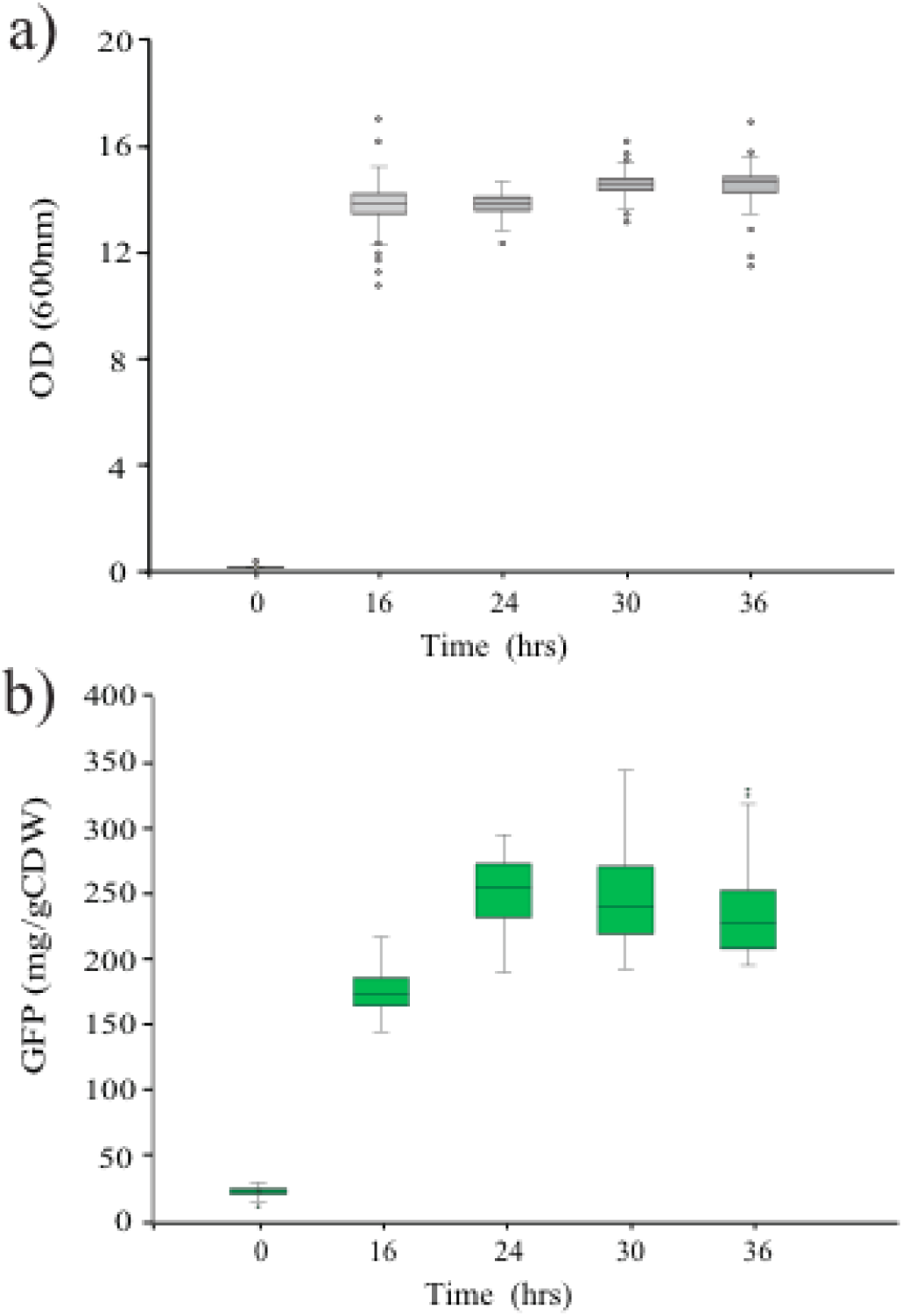
Autoinduced 2-stage microfermentatations. GFP and Biomass Levels throughout the Autoinduction Protocol (Figure 1a, Protocol 1). Biomass (a) and GFPuv levels (b) over time in AB2 media in microfermentations. (n=96)

**Figure 5:**
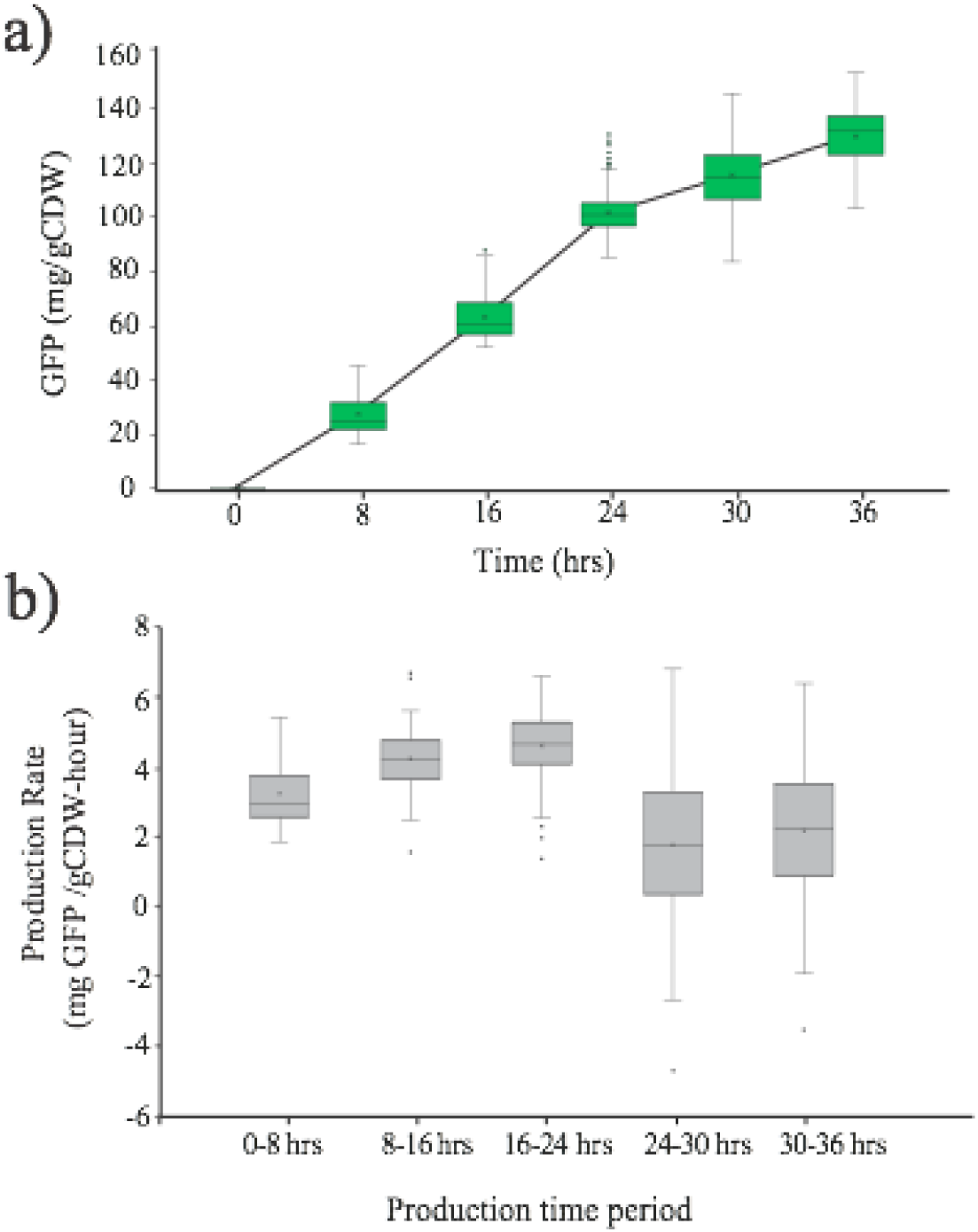
Wash based 2-stage microfermentations. a) Specific GFPuv production as a function of time, post-wash in the phosphate limited stationary phase. b) Specific GFPuv production rate as a function of time, post-wash in the phosphate limited stationary phase.

Additional specific details for the 2-stage wash protocol are also based on results given in Figure 6. The optimal incubation time in the SM10++ growth phase is 16 hrs (Figure 6a), which also conveniently enables inoculation at 5pm followed by harvest and washing the next morning at 9 am. As demonstrated in Figure 6b, changes in the biomass level used in the “production phase” will impact the final titer, however the specific production (titer/biomass level) is insensitive to these levels. As a result, we routinely measure optical densities at the end of the production phase and calculate specific production levels (g product/gram of biomass) or specific production rate (g product/gram of biomass - time). This reduces the requirement for a very accurate normalization step, which can be difficult in practice. It is also noteworthy, as illustrated in Figure 6c, that the cells do undergro one doubling (almost exactly) after washing to remove phosphate. Lastly, when performing the 2-stage wash protocol with numerous microtiter plates the time taken to wash and normalize is not insignificant and can take 1-2 hours if performing large screens. As a result, we evaluated the impact of the time cultures are held out of the incubator (at room temperature, without shaking) on final results. As can be seen in Figure 6d, reduced performance is observed only after holding the cultures over 2 hours out of the incubator. As a result, we recommend using a manageable number of plates to meet this constraint. Processing 10-20 microtiter plates should be manageable in 2 hrs, if more plates are required, these protocols are amenable to automation on liquid handling robots.

**Figure 6:**
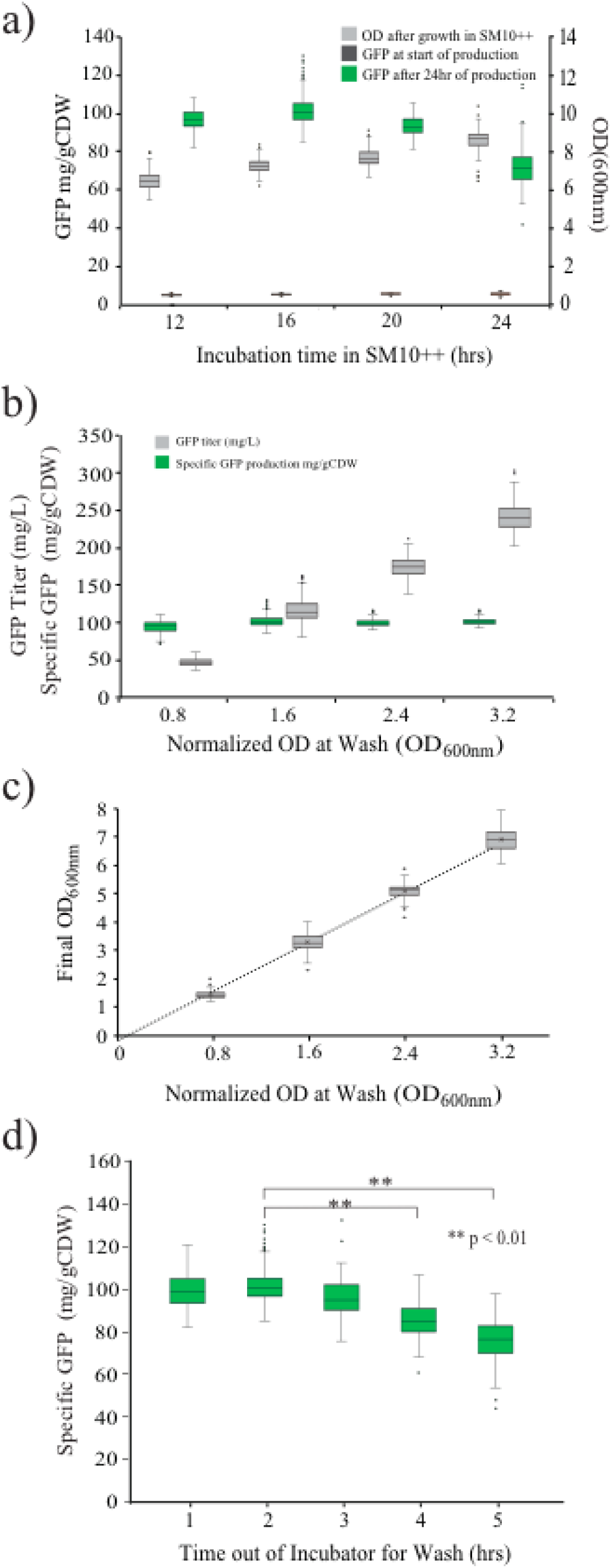
Impact of key protocol variables on output of the wash based 2-stage microfermentations. a) The impact of time spent in the growth phase (SM10++ culture, Protocol 2, Day 2) on final specific GFPuv expression levels at harvest. The impact of the normalized cell concentrations (optical densities) at the start of production on b) final GFPuv expression levels and specific production at harvest as well as c) final cell concentrations (biomass levels). d) The impact of the length of time cells are kept out of the incubator during the normalization step on final specific GFPuv expression levels at harvest. In these studies the phosphate free production phase was kept constant at 24 hrs.

While the current 2-stage wash protocol, as presented, has an optimal SM10++ “growth phase” of 16 hrs (Figure 6a), the length of the growth phase is not as critical as the stage of growth of the cells just prior to the wash step. Ideally, cells should be in mid exponential growth, just prior to wash, which in SM10++ media correlates with a culture OD_600nm_ from 5 to 15. Wells/cultures with low optical densities may still be in the lag phase, which can negatively affect performance in the production phase. If the wells/cultures have higher optical densities, they may well be entering stationary phase prior to the wash which can also negatively affect performance in the production phase. Of course this would be media dependent if adapting the protocol to another strain or media formulation. We recommend checking the optical density of cultures prior to the wash step and shortening or extending the growth phase as needed to ensure cultures are in mid exponential growth. Certain strains or strain variants may require a longer growth phase to reach mid exponential phase. We often encounter this when using strains with multiple plasmids requiring selection with multiple antibiotics.

### Applications

As discussed above these protocols can be leveraged to evaluate strains engineered to produce a wide variety of products from proteins and enzymes to small molecule chemicals. This approach can also be readily extended to enable expression of mutant protein libraries, where using strains additionally engineered for autolysis, enable rapid generation of cell lysates for subsequent screening. ^29^

### Comparison with other Methods

These protocols, as well as adaptations, offer a consistent approach to the standardized evaluation of microbial strains in batch cultures in microtiter plates. They offer a well outlined alternative to more complicated and expensive microbioreactor systems, and rely on inexpensive media, and consumables. The most expensive materials are the System Duetz, plate covers and clamps. This approach is readily scalable to experiments containing 10-20 microtiter plates enabling much higher throughput evaluations when compared to shake flasks, as well as most commercial and custom microbioreactor systems. Using standard laboratory automation (liquid handlers), the throughput can be greatly increased.

### Needed Expertise

There is no specialized expertise required to implement this protocol. Anyone with basic skills in microbiology and sterile technique should be able to successfully execute experiments utilizing these methods.

### Advantages

2-stage production has the potential to improve the performance, robustness and scalability of bioprocesses. ^22,44^ In addition and more specifically, dynamic metabolic control in the context of 2-stage production, enables optimization strategies that are not compatible with cellular growth, for example deletion of essential genes. ^45–47^ These microfermentation protocols also enable a high degree of automation, which in turn enables higher throughput strain evaluations at much lower costs than many commercial microbioreactor systems.

## Limitations

These protocols, as detailed, have developed based on *E. coli*, and the use of specifically engineered strains. The media components and culture/production environment have to be adjusted when applied to other microbial production hosts. We hope the approach to protocol development as discussed above, will enable adaptation of these methods to additional systems, including those with different microbial hosts, different induction systems (beyond phosphate limitation) and even 2-stage processes wherein the second stage is also a growth stage.

The microbial strain utilized for high throughput evaluation is also of critical importance. The *E. coli* strains presented in this protocol have been significantly engineered.^25,26,28,29^ Importantly, and somewhat uniquely these strains i) enable tightly controlled induction of promoters activated by phosphate depletion and ii) have no observable overflow metabolism in media with excess batch glucose.^25^ These traits enable reproducible 2-stage microfermentations. Firstly, tightly controlled expression is critical, if strains have significant leaky expression, this can affect lag times and growth rates leading to significant growth differences between strain variants being evaluated. This is not insurmountable but needs to be considered with the respect to the type of data being collected and how it is analyzed.

Overflow metabolism is a significant challenge in many strains of *E. coli*. As mentioned above, higher batch glucose levels enable higher biomass levels, as well as consequently high protein and small molecule titers. However many strains of *E. coli* produce overflow metabolites such as acetate when grown at high glucose levels. Acetate can accumulate, cause toxicity and affect production.^48^ One of the benefits of fed batch fermentations is the ability to keep residual glucose levels low to avoid acetate or overflow metabolite accumulation.^48^ Significant effort has been made to develop slow or time-released glucose formulations to effectively mimic fed batch conditions in small scale batch cultures. ^49–53^ To simplify batch microfermentations, strains with minimal overflow metabolism are ideal. If these protocols are adapted to other *E. coli* strains or microbial species, overflow metabolism should be considered when defining batch media, and establishing target biomass levels. Lower biomass levels can be achieved with lower batch glucose levels and should be considered if adapting to other strains of *E. coli*.

It is also worth noting that the data presented herein as well as previously reported results, leveraged promoters induced upon phosphate depletion that are, importantly, also robust. Expression levels are consistent across production scales and media formulations. ^24^

### Experimental Design

These protocols, as well as any adaptations, will often comprise the evaluation or testing part of a design, build, test, learn (DBTL) cycle. ^54,55^ As a result critical aspects of the final experimental design need to account for other components of these cycles. However, we do recommend running standard internal process controls on each microtiter plate. These controls for example could include a strain expressing GFP or mCherry (or another easily measured product). This enables tracking of “process” performance between plates over time. A control chart can then be developed, and plates whose internal controls fall outside of an acceptable range can be repeated. This is best practice with any high throughput evaluation. ^43^

## Materials

### Consumables & Reagents

1. *E. coli* strain DLF_R004, Roke Biotechnologies, LLC., Catalog # RSA-002. Genotype: F-, λ-, Δ(araD-araB)567, lacZ4787(del)(::rrnB-3), rph-1, Δ(rhaD-rhaB)568, hsdR51, ΔackA-pta, ΔpoxB, ΔpflB, ΔldhA, ΔadhE, ΔiclR, ΔarcA ΔompT yibDp-λR-nucA::apmR
2. *E. coli* strain DLFS_0025 Genotype: F-, λ-, Δ(araD-araB)567, lacZ4787(del)(::rrnB-3), rph-1, Δ(rhaD-rhaB)568, hsdR514, ΔackA-pta, ΔpoxB, ΔpflB, ΔldhA, ΔadhE, ΔiclR, ΔarcA, ΔsspB::frt, Δcas3:: ugpBp-sspB-yibDp-casA
3. Plasmid pHCKan-GFPuv, Addgene #127078
4. Sterile Flat Bottom 96 well plates, Genesee Scientific, Catalog # 25-104
5. Sterile U Bottom 96 well plates, Genesee Scientific, Catalog # 25-224
6. System Duetz Covers: Enzyscreen, Part #CR1596
7. System Duetz Clamps: Enzyscreen, Part #CR1800
8. Multi-Channel Pipette(s) capable of transferring volumes from 10-200 µL
9. Black-walled 96-well plates for measuring Fluorescence (655087, Greiner Bio-One)
10. High mass transfer FlowerPlate (Cat #MTP-48-B; m2p-labs)
11. Sterile Syringe Filters, Genesee Scientific, Catalog # 25-244
12. Sterile Vacuum-Driven Filter, Genesee Scientific, Catalog # 25-233

### Media Components

1. LB Lennox Formulation 10 g/L tryptone, 5 g/Lyeast extract, and 5g/L sodium chloride per liter.
2. AB2 Media For a detailed preparation process refer to Menacho-Melgar et al ^23^. Components: 6.2 g/L yeast extract, 3.5 g/L casamino acids, 5.4 g/L ammonium sulfate anhydrous and 41.8 g/L Bis-Tris, 45g/l glucose. pH is adjusted to 6.8.
3. SM10 ++ Media (Refer to Supplemental Materials)
4. SM10 No Phosphate Media (Refer to Supplemental Materials)

### Equipment

1. Shaking incubator capable of temperatures of 37 degree Celius, shaking speeds of 300 rpm and a shaking orbit of 50 mm. Example: Kuhner: Climo-Shaker ISF4-X
2. A centrifuge capable of handling microtiter plates at g forces of 3500rpm (2900 rcf). Example: Thermo Sorvall Legend XTR Refrigerated Centrifuge
3. Microtiter Absorbance Plate Reader: Tecan Infinite 200 or Molecular Devices Example: Spectramax Plus 384 Benchtop Cuvette Plate Spectrophotometer
4. Microtiter Fluorescence Plate Reader : Example: Tecan Infinite 200
5. Fermentation monitoring system. Example: M2P labs: BioLector® I, 48 Parallel Microbioreactors.

#### Protocol 1: 2-stage autoinduction microfermentation protocol

### Day 1 (Starter Cultures)

Timing: ∼ 10 min per microtiter plate

1. Start LB cultures in sterile flat bottom 96 well microtiter plates. The fill volume for each well should be 150 *µ*L of LB, Lennox formulation plus 35 ug/mL kanamycin, or alternative antibiotics as appropriate. Single colonies, previously LB cultures or cryostocks can be used for inoculation. When using prior cultures or cryostocks we recommend inoculating 145 *µ*L of media with 5ul of culture/cryostock. Preferably, do not utilize wells on the outer edge of the plate.
2. Cover the 96 well plates with sterilized plate covers from Enzyscreen and secure in a shaking incubator with appropriate clamps.
3. Incubate at 37°C for 16 hrs, shaking at 300 rpm, with a 50 mm shaking orbit.

### Day 2 (Autoinduction Cultures)

Timing: ∼ 10 min per microtiter plate

1. For each LB starter plate, fill a fresh sterile flat bottom 96 well microtiter plate with 145 *µ*L of AB2 media plus appropriate antibiotics. AB Media may be used in lieu of AB2.
2. Inoculate the AB2 plates with 5ul of the LB starter cultures.
3. Cover the new AB2 with a fresh sterile plate cover and secure in a shaking incubator with appropriate clamps.
4. Incubate at 37 °C for 24 hrs, shaking at 300 rpm, with a 50 mm shaking orbit.

### Day 3 (Harvest)

Timing: ∼ 30 min per microtiter plate

1. For each AB2 culture plate, obtain 1 new flat bottom 96 well plate, 1 black-walled 96-well plate and 1 U-bottom 96 well plate.
2. Fill the wells of one flat bottom 96 well plate with 190 *µ*L of deionized water (OD plate).
3. To each well of the OD plate, transfer 10 *µ*L of AB2 culture from the AB2 culture plate to generate a 20 fold diluted culture for measuring optical densities.
4. Measure the optical density of the OD plate at 600 nm using an Absorbance Plate Reader.
5. Calculate the harvest optical densities for each well by correcting the raw readings, according to Equation 1. The path length correction factor should be measured for each plate type and plate reader. Using the equipment and plates listed in the protocol the path length correction is 1.6

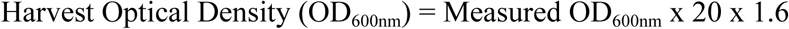
6. The 20X diluted sample, or further diluted samples, can also be used directly for final analyses requiring suspended cells, such as relative fluorescence of GFP. In these studies, for GFPuv, fluorescence measurement was performed using the black-walled 96-well plates. The diluted samples were excited at 412 nm (bandwidth 10nm) and then we measured the emission at 530 nm (bandwidth=10nm). And the gain was set at 60. The corresponding coefficient between the GFPuv fluorescence units and the mass of GFPuv is 3.24 e9, meaning 1 g of GFPuv will be corresponded to 3.24 e9 relative fluorescent units. ^23^ Then the relative GFP production will be normalized by the cell biomass (OD600nm x 0.35).

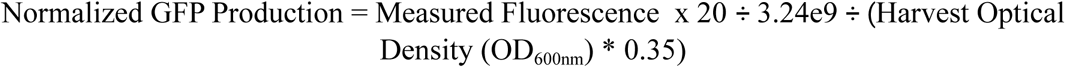
7. Transfer ther remainder of the AB2 culture (∼ 75 *µ*L) from each well to the U bottom 96 well plate. Due to evaporation, the final volume of the AB2 cultures will be ∼ 130uL, and after taking 20 *µ*L to measure the optical density, we recommend transferring 75 *µ*L.
8. Centrifuge the U-bottom 96 well plates at 3500 rcf for 10 minutes.
9. After centrifugation, carefully transfer the supernatant from each well to the second fresh flat bottom 96 well plate. Be careful not to disturb the pellet. If the pellet is to be discarded we recommend transferring 60 *µ*L. If the pellet is to be analyzed we recommend carefully aspirating as much supernatant as possible.
10. The supernatant and cell pellets are now ready for subsequent analyses.

#### Protocol 2: 2-stage wash microfermentation protocol

### Day 1 (Starter Cultures)

Timing: ∼ 10 min per microtiter plate

1. Start LB cultures in sterile flat bottom 96 well microtiter plates. The fill volume for each well should be 150 *µ*L of LB, Lennox formulation plus 35 ug/mL kanamycin, or alternative antibiotics as appropriate. Single colonies, previously LB cultures or cryostocks can be used for inoculation. When using prior cultures or cryostocks we recommend inoculating 145 *µ*L of media with 5ul of culture/cryostock. Preferably, do not utilize wells on the outer edge of the plate.
2. Cover the 96 well plates with sterilized plate covers from Enzyscreen and secure in a shaking incubator with appropriate clamps.
3. Incubate at 37 °C for 16 hrs, shaking at 300 rpm, with a 50 mm shaking orbit.

### Day 2 (SM10++ Growth Cultures)

Timing: ∼ 10 min per microtiter plate

11. For each LB starter plate, fill a fresh sterile flat bottom 96 well microtiter plate with 145 *µ*L of SM10++ media plus appropriate antibiotics.
12. Inoculate the SM10++ plates with 5*µ*L of the LB starter cultures.
13. Cover the new SM10++ with a fresh sterile plate cover and secure in a shaking incubator with appropriate clamps.
4. Incubate at 37 °C for 16 hrs, shaking at 300 rpm, with a 50 mm shaking orbit.

### Day 3 (Wash and Normalization)

Timing: ∼ 30 min per microtiter plate

1. For each SM10++ culture plate, obtain 2 new flat bottom 96 well plates and 1 U-bottom 96 well plate.
2. Fill the wells of one flat bottom 96 well plate with 190 *µ*L of deionized water (OD plate).
3. To each well of the OD plate, transfer 10 *µ*L of SM10++ culture from the SM10++ culture plate to generate a 20 fold diluted culture for measuring optical densities.
4. Measure the optical density of the OD plate at 600 nm using an Absorbance Plate Reader.
5. Calculate the harvest optical densities for each well by correcting the raw readings, according to Equation 1. The path length correction factor should be measured for each plate type and plate reader. Using he equipment and plates listed in the protocol the path length correction is 1.6

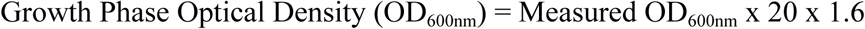 CRITICAL: For each 96 well plate optical densities should range between 5-15. Wells outside this range should be noted during subsequent analysis.
6. Transfer ther remainder of the SM10++ culture (∼ 120 *µ*L) from each well to the U bottom 96 well plate. Due to evaporation, the final volume of the AB2 cultures will be ∼ 130 *µ*L, and after taking 10 *µ*L to measure the optical density, we recommend transferring 120 *µ*L.
7. Centrifuge the U-bottom 96 well plates at 3500 rpm (2900 rcf) for 10 minutes.
8. After centrifugation, carefully aspirate and discard all of the supernatant from each well. Be careful not to disturb the pellet. CRITICAL: It is important to remove all supernatant and remaining phosphate, even if some cells are lost.
9. To each well add 140 *µ*L of SM10 No Phosphate Media and resuspend the pellets in the SM10 No Phosphate Media by pipetting up and down.Centrifuge the U-bottom 96 well plates again at 3500 rpm (2900 rcf) for 10 minutes.
10. Take 90 *µ*L of supernatant out and there will be about 50 *µ*L left in each well. Then mix the left pellet and supernatant by pipetting up and down.
11. Measure the OD600nm of the concentrated cells using the same way as above, 20 x dilution and then plate reader.
12. Calculate the volume needed for the target normalized OD in a total 150 *µ*L volume.
13. In the case of GFPuv, fill the wells of the second flat bottom 96 well plate with 133 *µ*L of SM10 No Phosphate Media, plus appropriate antibiotics. Then transfer 17 *µ*L of resuspended culture to the appropriate wells
14. Cover the new SM10 No Phosphate Plate with a fresh sterile plate cover and secure in a shaking incubator with appropriate clamps.
15. Incubate at 37 °C for 24 hrs, shaking at 300 rpm, with a 50 mm shaking orbit.

### Day 4 (Harvest)

Timing: ∼ 30 min per microtiter plate

1. For each SM10 No Phosphate culture plate, obtain 1 new flat bottom 96 well plate, 1 black-walled 96-well plate and 1 U-bottom 96 well plate.
2. Fill the wells of one flat bottom 96 well plate with 190 *µ*L of deionized water (OD plate).
3. To each well of the OD plate, transfer 10 *µ*L of SM10 No Phosphate culture from the SM10 No Phosphate culture plate to generate a 20 fold diluted culture for measuring optical densities.
4. Measure the optical density of the OD plate at 600 nm using an Absorbance Plate Reader.
5. Calculate the harvest optical densities for each well by correcting the raw readings, according to Equation 1. The path length correction factor should be measured for each plate type and plate reader. Using he equipment and plates listed in the protocol the path length correction is 1.6

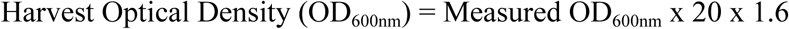 CRITICAL: When comparing variants/well, product titers should be normalized by the Harvest Optical Density for each well.
5. The 20X diluted sample, or further diluted samples, can also be used directly for final analyses requiring suspended cells, such as relative fluorescence of GFP. In these studies, for GFPuv, fluorescence measurement was performed using the black-walled 96-well plates.

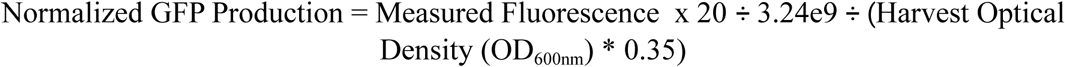
6. Transfer ther remainder of the SM10 No Phosphate culture (∼ 75 *µ*L) from each well to the U bottom 96 well plate. Due to evaporation, the final volume of the AB2 cultures will be ∼ 100 *µ*L, and after taking 20 *µ*L to measure the optical density, we recommend transferring 75 *µ*L.
7. Centrifuge the U-bottom 96 well plates at 3500 rpm (2900 rcf) for 10 minutes.
8. After centrifugation, carefully transfer the supernatant from each well to the second fresh flat bottom 96 well plate. Be careful not to disturb the pellet. If the pellet is to be discarded we recommend transferring 60 *µ*L. If the pellet is to be analyzed we recommend carefully aspirating as much supernatant as possible.
9. The supernatant and cell pellets are now ready for subsequent analyses.

## Conflict of interests

RMM and MDL have financial interests in Roke Biotechnologies, LLC. MDL and YE have a financial interest in DMC Biotechnologies, Inc.

## Author Contributions

SL, ZY, EAM, RMM and MDL conceived the study, performed experiments, prepared and edited the manuscript.

## Acknowledgements

We would like to acknowledge the following support: the NSF EAGER: #1445726 DARPA# HR0011-14-C-0075, as well as the North Carolina Biotechnology Center 2018-BIG-6503. R. Menacho-Melgar was supported in part by the NIH Biotechnology Training Grant (T32GM008555).

