## Supplementary Materials for "The Development of 2-stage Microfermentation Protocols for High Throughput Cell Factory Evaluations"

### Media Preparation Protocols

#### Materials used for stock solutions

Stock solutions were prepared using ultrapure water and ACS grade reagents

| Name | Supplier | Catalog/Item Number |
|---|---|---|
| Tryptone Powder | Biobasic | TG217(G211) |
| Yeast Extract | Biobasic | G0961 |
| Sodium Chloride | Biobasic | DB0483 |
| Casamino Acid | Biobasic | CB3060 |
| Ammonium sulfate | Biobasic | ADB0060 |
| Citric Acid | Alfa Aesar | 36664 |
| Ferrous sulfate heptahydrate | Sigma | 15422-250G |
| Potassium phosphate dibasic | Sigma | P3786-1KG |
| Potassium phosphate monobasic | Sigma | P5655-1KG |
| $CaSO_4 \cdot 2H_2O$ | Alfa Aesar | 36700 |
| $ZnSO_4 \cdot H_2O$ | Sigma | 307491-100G |
| $MnSO_4 \cdot H_2O$ | Alfa Aesar | 33341 |
| $CuSO_4 \cdot 5H_2O$ | Alfa Aesar | 14178 |

| | | |
|---|---|---|
| $CoSO_4 \cdot 7H_2O$ | Acros Organics | AC213105000 |
| $MoNa_2O_4$*$2H_2O$ | Alfa Aesar | 12214 |
| Potassium 3-(N-morpholino) propane sulfonic acid (MOPS) | Biobasic | MB0360 |
| $H_3BO_3$ | Spectrum | B1125 |
| Thiamine Hydrochloride | Sigma | T4625-25G |
| Magnesium sulfate heptahydrate | Sigma | 230391-1KG |

##### LB media

For 1 liter of LB media, 10 g of tryptone, 5 g of yeast extract, and 5g of sodium chloride were added to 1 liter of ultrapure water. Then, autoclave the mixture to sterilize it.

##### AB2 media

Part 1: 5.4 g $(NH_4)_2SO_4$, 3.5 g Casamino acids, 41.8 g of Bis-Tris and 6.2 g yeast extract were mixed and dissolved in 910 ml of water and 3 ml of 12 M HCl. autoclaved in 910 ml of water and 3 ml of 12 M HCl. Then the mixture was autoclaved. Part 2: 500 g/L glucose was prepared and autoclaved. Finally, 910 mL of part 1 and 90 mL of part 2 were mixed.

##### SM10 and SM10++ media

**Step 1: stock solution preparation**

A. Ammonium-Citrate salts: Prepare 1 liter of 10X concentrated Ammonium-Citrate 90 salts by mixing 90 g of $(NH_4)_2SO_4$ and 2.5 g Citric Acid in water with stirring. Autoclave and store at RT.
B. Buffering reagents:
   a. Prepare 1 M Potassium 3-(N-morpholino) propane sulfonic acid (MOPS) and adjust to pH 7.4 with KOH (~40 mL). Filter sterilize (0.2 um) and store at RT.
   b. Prepare 0.5 M potassium phosphate buffer, pH 6.8 by mixing 248.5 mL of 1.0 M $K_2HPO_4$ and 251.5 mL of 1.0 M $KH_2PO_4$ and adjust to a final volume of 1000 mL with ultrapure water. Filter sterilize (0.2 um) and store at RT.
C. Prepare 2 M $MgSO_4$ and 10 mM $CaSO_4$ solutions. Filter sterilize (0.2 um) and store at RT.
D. Prepare a solution of micronutrients in 1000 mL of water containing 10 mL of concentrated $H_2SO_4$: 0.6 g $CoSO_4$*$7H_20$, 5.0 g $CuSO_4$*$5H_20$, 0.6 g $ZnSO_4$*$7H_20$, 0.2 g $MoNa_2O_4$*$2H_2O$, 0.1 g $H_3BO_3$, and 0.3 g $MnSO_4$*$H_2O$. Filter sterilize (0.2 um) and store at RT in the dark.
E. Prepare a fresh solution of 40 mM ferric sulfate heptahydrate in water. Filter sterilize (0.2 um) and discard after 1 day.
F. Prepare a 50 g/L solution of thiamine-HCl. Filter sterilize (0.2 um) and store at 4°C.

G. Prepare a 500 g/L solution of glucose by stirring with heat. Cool, filter sterilize (0.2 um), and store at RT.

**Step 2: Media preparation**

Different media were prepared according to the formulas described in the tables below.

| **SM10 ++ (pH 6.8)** | | | |
|---|---|---|---|
| **Ingredient** | **Concentration Stock** | **Volume in 1 L (mL)** | **Final Concentration** |
| **Ammonium-Citrate 90 Salts, pH 7.5** | 10 X FGM10 Salt (90 g Ammonium Sulfate, 2.5 g Citrate) | 100.0 | 1 X (9 g Ammonium Sulfate, 0.25 g Citrate) |
| **Phosphate Buffer, pH 6.8** | 500 mM | 10.0 | 5.00 mM |
| **Trace Metals** | 500 X | 4.0 | 2 X |
| **Fe (II) Sulfate** | 40 mM | 4.0 | 0.16 mM |
| **MgSO4** | 2 M | 1.25 | 2.50 mM |
| **CaSO4** | 10 mM | 6.25 | 0.06 mM |
| **Glucose** | 500 g/L | 90.0 | 45 .0g/L |
| **MOPS** | 1 M | 200.0 | 200 mM |
| **Thiamine-HCl** | 50 g/L | 0.2 | 0.01 g/L |
| **Yeast Extract** | 100 g/L | 25.0 | 2.5 g/L |
| **Casamino Acids** | 100 g/L | 25.0 | 2.5 g/L |

| **SM10 No Phosphate (pH 6.8)** | | | |
|---|---|---|---|
| **Ingredient** | **Concentration Stock** | **Volume in 1 L (mL)** | **Final Concentration** |
| **Ammonium-Citrate 90 Salts, pH 7.5** | 10 X FGM10 Salt (90 g Ammonium Sulfate, 2.5 g Citrate) | 100.0 | 1 X (9 g Ammonium Sulfate, 0.25 g Citrate) |

| | | | |
|---|---|---|---|
| **Trace Metals** | 500 X | 4.0 | 2 X |
| **Fe (II) Sulfate** | 40 mM | 4.0 | 0.16 mM |
| **MgSO4** | 2 M | 1.25 | 2.50 mM |
| **CaSO4** | 10 mM | 6.25 | 0.06 mM |
| **Glucose** | 500 g/L | 90.0 | 45 .0g/L |
| **MOPS** | 1 M | 200.0 | 200 mM |
| **Thiamine-HCl** | 50 g/L | 0.2 | 0.01 g/L |
